# Dissociable roles of human frontal eye fields and early visual cortex in presaccadic attention – evidence from TMS

**DOI:** 10.1101/2023.02.23.529691

**Authors:** Nina M. Hanning, Antonio Fernández, Marisa Carrasco

## Abstract

Shortly before each saccadic eye movement, presaccadic attention improves visual sensitivity at the saccade target^1–5^ at the expense of lowered sensitivity at non-target locations^6–11^. Some behavioral and neural correlates of presaccadic attention and covert attention –which likewise enhances sensitivity, but during fixation^12^–are similar^13^. This resemblance has led to the debatable^13–18^ notion that presaccadic and covert attention are functionally equivalent and rely on the same neural circuitry^19–21^. At a broad scale, oculomotor brain structures (e.g., FEF) are also modulated during covert attention^22–24^ – yet by distinct neuronal subpopulations^25–28^. Perceptual benefits of presaccadic attention rely on feedback from oculomotor structures to visual cortices^29,30^ (**Fig. 1a**); micro-stimulation of FEF in non-human primates affects activity in visual cortex^31–34^ and enhances visual sensitivity at the movement field of the stimulated neurons^35–37^. Similar feedback projections seem to exist in humans: FEF+ activation precedes occipital activation during saccade preparation^38,39^ and FEF TMS modulates activity in visual cortex^40–42^ and enhances perceived contrast in the contralateral hemifield^40^. We investigated presaccadic feedback in humans by applying TMS to frontal or visual areas during saccade preparation. By simultaneously measuring perceptual performance, we show the causal and differential roles of these brain regions in contralateral presaccadic benefits at the saccade target and costs at non-targets: Whereas rFEF+ stimulation reduced presaccadic costs throughout saccade preparation, V1/V2 stimulation reduced benefits only shortly before saccade onset. These effects provide causal evidence that presaccadic attention modulates perception through cortico-cortical feedback and further dissociate presaccadic and covert attention.

## Introduction

Every time we open our eyes, we confront an overwhelming amount of visual information, yet we have the impression of effortlessly understanding what we see. We are typically not aware of the complex neuro-cognitive processes that help us prioritize the visual input. Visual attention allows us to selectively filter relevant information out of irrelevant noise by prioritizing some aspects of the visual scene while ignoring others^12,43^. This attentional selection is usually achieved by a succession of rapid saccadic eye movements toward relevant information of the visual scene^44^. Interestingly, attention reaches the next relevant location already before the eyes start to move. This *presaccadic shift of attention* is indicated by perceptual benefits at the saccade target^1–5^, at the expense of costs elsewhere^6–11^.

Presaccadic attention modulates the contrast response function (CRF), which characterizes the nonlinear, sigmoidal relation between the contrast (or intensity) of a visual stimulus and the resulting response^45–47^, such as neuronal firing rate or, consequently, visual performance. Specifically, presaccadic attention alters perceptual performance via response gain, i.e., it increases d_max_ (the maximal response at high contrast) at the saccade target (**Fig. 1b**), and reduces d_max_ at non-target locations^9,48^.

**Figure 1.**
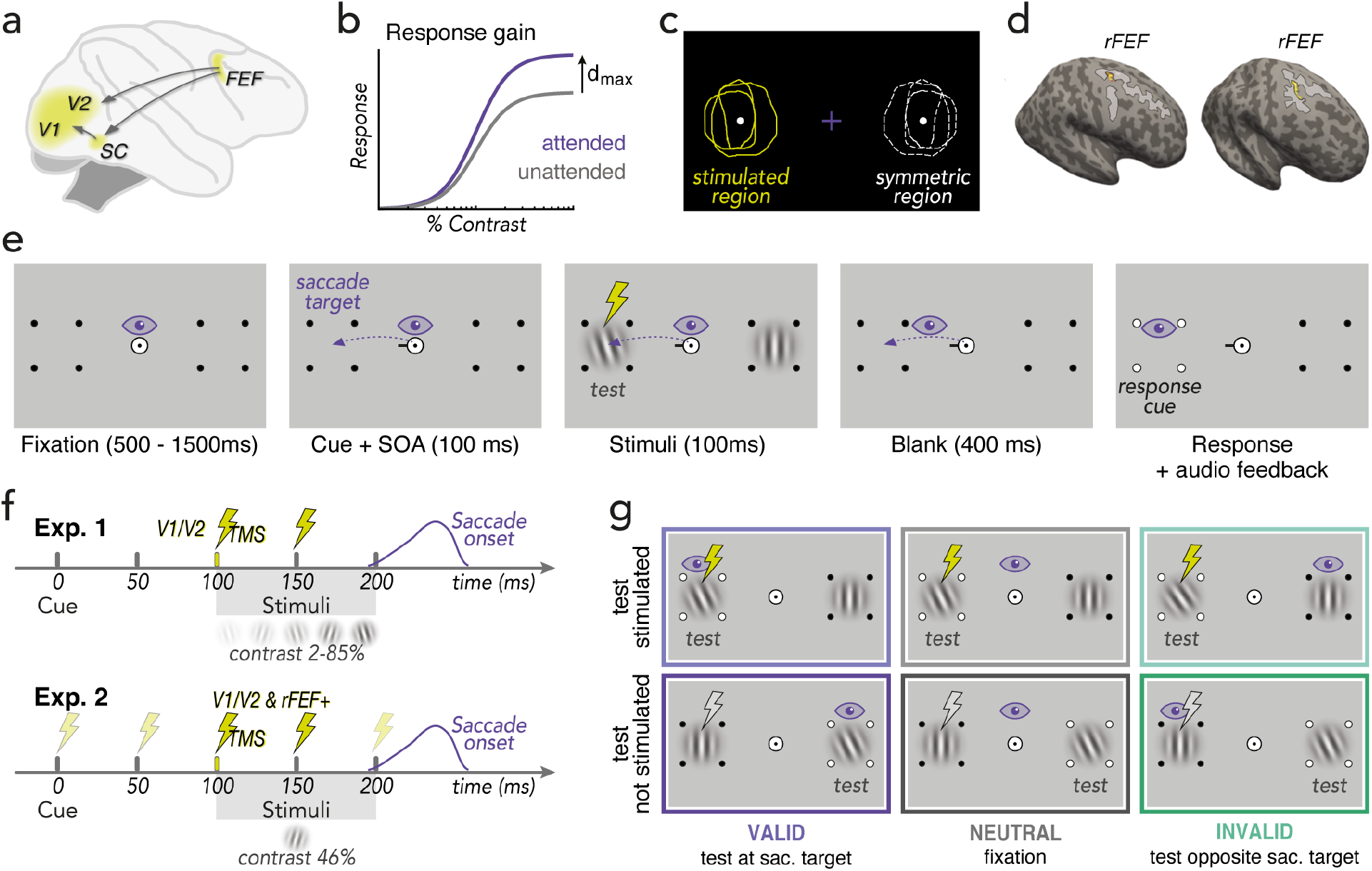
Background and experimental design. (**a**) Feedback connections originally established in non-human primates^29,30,36,90,104,105^ assumed to underly perceptual correlates of human presaccadic attention. *FEF:* Frontal eye field, *SC*: Superior Colliculus; (**b**) Response gain effect of presaccadic attention on the contrast response function: presaccadic attention scales the response by a multiplicative gain factor, resulting in an increase of the asymptotic response d_max_ (the maximal response achieved at high contrast)^9,48^. (**c**) Determining occipital (V1/V2) stimulation sites (Exp.1 & 2a) and stimulus placement for both experiments via ‘phosphene mapping’: Observers were stimulated laterally around the occipital pole until they perceived a phosphene (in the contralateral visual field), then drew its outline on the screen. The center of the phosphene drawings (*stimulated region*) and the *symmetric region* (not stimulated) were used for stimulus placement in the main experiment, where we applied sub-phosphene-threshold TMS using identical coil positioning. (**d**) Determining rFEF+ stimulation sites (**Exp.2b**). rFEF+ (yellow ROI) was localized on each individual observer’s anatomy (two exemplary observers shown here) via a probabilistic topography atlas^76^ and verified by anatomical landmarks (junction of the precentral and superior frontal sulcus; light gray areas)^64,65,67^. (**e**) Presaccadic orientation discrimination task. After a fixation period, a central direction cue (black line) appeared. Observers were instructed to make a saccade to the indicated target marked by placeholder dots. Note that the saccade was equally likely directed to the stimulated and to the symmetric region / hemifield. 100ms after cue onset, a tilted test Gabor patch was presented at either the saccade target (*valid; 50%*) or at the opposite location (*invalid*), randomly intermixed; a vertical Gabor was presented at the other location. Importantly, stimuli were presented during saccade preparation, i.e., while gaze was still at the screen center. After saccade offset, a response cue (white dots on placeholder) indicated the location at which the test Gabor had appeared, and observers reported its orientation. In the *neutral* condition (separately blocked) the cue pointed to both placeholders and observers kept fixating (supplemental **Video S1** demonstrates the trials sequence of each condition). (**f**) Trial timeline. Observers received double-pulse TMS (50ms inter-pulse interval) locked to stimuli onset (Exp.1) or at various times during saccade preparation (0-200ms relative to cue onset; Exp.2). Whereas grating contrast was varied to measure contrast response functions in Exp.1, contrast was fixed to 46% in Exp.2. (**g**) Experimental conditions. The test was equally likely presented at the stimulated region (*test stimulated*) or in the opposite hemifield (*not stimulated*). Moreover, the test was equally likely presented at the saccade target (*valid*) or opposite of it (*invalid*). Note that the fixation condition (*neutral*) was only tested in Exp.1.

This characteristic ‘push-pull mechanism’ is also observed for *covert* attentional orienting, in the absence of eye movements: Behaviorally relevant items, to which we voluntarily deploy *endogenous* attention, as well as salient events that automatically capture *exogenous* attention, likewise cause perceptual benefits at the attended and concomitant costs at unattended locations^9,12,49,50^. Both covert and presaccadic attention modulate visual processing along the cortical hierarchy. Merely shifting the focus of attention (while keeping the retinal image constant) affects neuronal responses^26,51–55^. As the perceptual and neuronal dynamics of covert attention seemingly mimic those of saccade preparation, some have postulated that the same neural mechanism underlies the two processes; even covert shifts of attention (during fixation) would result from eye movement planning^19–21^.

Human neuroimaging studies indicate that saccade planning and covert exogenous and endogenous attention differentially modulate brain activity^55–58^. Yet, these studies cannot establish causality, as fMRI techniques record, but do not manipulate brain function. Using transcranial magnetic stimulation (TMS) to briefly, and non-invasively, alter cortical activity^59–62^, we recently dissociated the causal role of two brain areas involved in covert exogenous and endogenous: Whereas occipital stimulation extinguished benefits and costs of exogenous attention^50^, it did not alter endogenous attention^63^ – which instead was affected by stimulation of rFEF+^63^, the human homolog of the right macaque frontal eye field^64–67^. Saccade preparation enhances neural responses in oculomotor areas and elicits retinotopic activity in visual cortex^55,68^, but it is unknown which brain areas play a *causal* role in presaccadic attention.

The aim of the present study was two-fold: (*1*) test and potentially dissociate the causal involvement of early visual areas (V1/V2) in presaccadic attention dynamics from those recently established for covert attention^50,63^; (*2*) elucidate the differential role of and interplay between frontal (rFEF+) and visual areas (V1/V2) in perceptual benefits and costs preceding saccadic eye movements to investigate the role of cortico-cortical feedback in presaccadic attention.

## Results

To assess whether early visual areas are critical for presaccadic attention, **in Experiment 1** we measured CRFs during fixation or saccade preparation in a combined psychophysics-TMS experiment. Saccades were prepared either to the visual field location affected by V1/V2 TMS, individually determined using an established phosphene mapping procedure^50,63,69,70^ (**Fig. 1c**; *Methods – Phosphene mapping & stimulus placement*), or to the symmetric region in the other hemifield (internal control, unaffected by TMS). Shortly before saccade onset, we briefly presented oriented test stimuli either at or opposite to the saccade target (**Fig. 1e**) and applied (sub-threshold) double-pulse TMS to the individually determined V1/V2 stimulation site (**Fig. 1f**). We tested multiple stimulus contrasts and derived CRFs for all combinations of presaccadic attention at the stimulated and non-stimulated symmetric location (**Fig. 1g**), to characterize the effect of TMS on performance benefits and costs relative to fixation.

### Differential role of early visual cortex in presaccadic and covert attention (Exp.1)

To assess whether TMS to early visual areas affects presaccadic attentional modulations, we obtained CRFs by fitting Naka-Rushton functions^45^ to the visual sensitivity data (**Fig. 2a**; *Methods – Quantification & statistical analysis*). To evaluate the predicted response gain effect of presaccadic attention on the upper asymptote d_max_^9,48^ as well as a potential effect of V1/V2 TMS, we conducted a repeated measures ANOVA [presaccadic attention (valid/neutral/invalid) * TMS_site (test stimulated/not stimulated)]. Presaccadic attention modulated the upper asymptote of the functions (*F*(2,18)=164.34, *p*<0.001), which, compared to *neutral* (fixation baseline), was significantly increased at the saccade target (*valid*; *p*<0.001, *d*=1.82) and decreased at the non-target (*invalid*; *p*<0.001, *d*=5.26) – reflecting the typical presaccadic benefit and cost, respectively. Importantly, TMS site neither affected asymptotic performance (*F*(1,9)<1), nor did it interact with presaccadic attention (*F*(2,18)<1).

**Figure 2.**
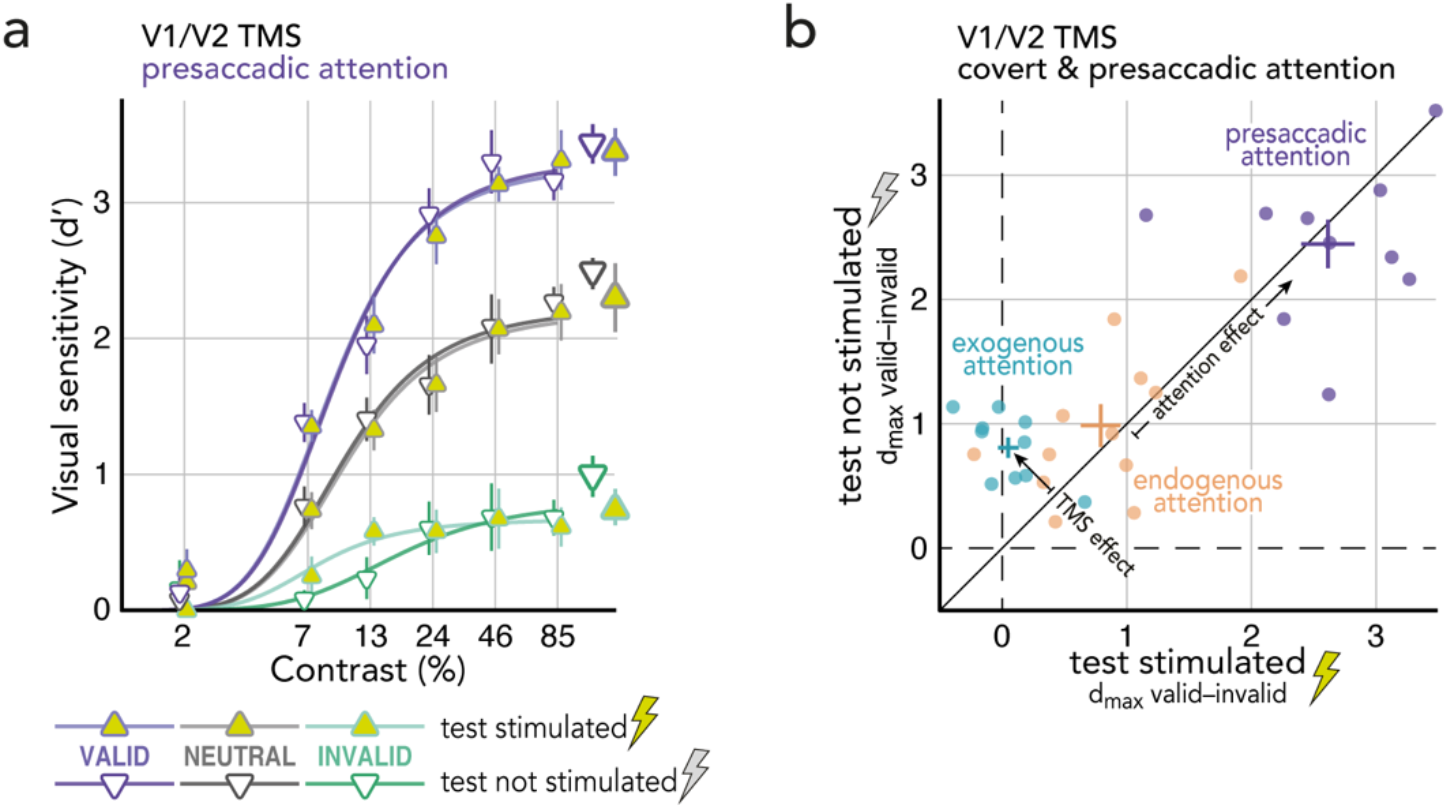
The effect of V1/V2 TMS on presaccadic attention. (**a**) Contrast Response Functions (CRF) at the saccade target (purple), opposite the saccade target (green), or during fixation (gray; baseline), measured at the stimulated region (yellow symbols) or in the non-stimulated hemifield (white symbols). Respective group averaged parameter estimates for the upper asymptote d_max_ (based individual observers’ fits) displayed on the right. Error bars indicate ±1SEM. (**b**) The effect of V1/V2 TMS on presaccadic (purple) and covert exogenous (blue) and endogenous attention (orange). Dots represent individual observers’ attention effects (valid minus invalid d_max_ estimates) at the stimulated (x-axis) plotted against the not stimulated region (y-axis). Crosses represent the group mean ±1SEM.

This finding is in direct contrast to the effect of covert exogenous attention^50^: In the same psychophysics-TMS protocol (except that instead of a central saccade cue, a peripheral attention cue modulated attention), V1/2 TMS eliminated response gain benefits and costs of exogenous attention (**Fig. 2b**, exogenous attention & **Fig. S1a**). Covert endogenous attention (modulated by a central cue indicating the likely test location), however, like presaccadic attention, was not affected by V1/V2 TMS^63^ (**Fig. 2b**, endogenous attention & **Fig. S1b**). These results reveal a neural dissociation of presaccadic and covert exogenous attention. Moreover, they show that the perceptual effects of presaccadic attention (i.e., the difference between *valid* and *invalid*) are stronger than those of covert exogenous and endogenous attention (**Fig. 2b**). This also applies to the separate benefits (difference *valid* and *neutral*) and costs (difference *invalid* and *neutral*), which are both more pronounced for presaccadic attention (**Fig. 2a**) than for covert exogenous (**Fig. S1a**) and endogenous (**Fig. S1b**) attention.

When investigating the role of early visual areas in covert attention^50,63^, we applied V1/V2 TMS at the known peak of exogenous and endogenous attention effects on performance^12,71^, i.e., 100ms^50^ and 500ms^63^ after the respective cue onset. Presaccadic attention builds up during saccade preparation, gradually reaching its maximum shortly before saccade onset – which typically is ~200ms after saccade cue onset^4–6,72–75^, but this varies with the pronounced variance in saccade latencies among and within individual observers.

To investigate the role of early visual areas throughout saccade preparation (including the time of peak presaccadic attention right before saccade onset), in **Experiment 2a** we stimulated V1/V2 at various timepoints after saccade cue onset (**Fig. 1f**), using the otherwise identical psychophysics-TMS protocol. Given the assumption that feedback from higher-order to early visual areas underlies presaccadic attention modulations, V1/V2 should contribute to (and thus V1/V2 TMS should affect) presaccadic attention at later stages of saccade programming, whereas TMS of the frontal eye fields (FEF) – an area providing the source of feedback, which we stimulated in **Experiment 2b** – should show a relatively earlier effect. V1/V2 stimulation site and stimulus placement were determined via phosphene mapping^50,63,69,70^ (**Fig. 1c**). We localized rFEF+, the putative human homolog of macaque frontal eye field^64–67^, on each observer’s anatomical brain scan (acquired via MRI) via a probabilistic topography atlas^76^ (**Fig. 1d**; *Methods – FEF Localization*).

For a temporal analysis of the effect of V1/V2 and FEF+ TMS on saccade latencies and landing precision, see supplemental **Fig. S2**. Overall, presaccadic TMS slowed down saccadic reaction times: The later the double-pulse was applied (after the saccade cue), the longer the saccade latencies. This effect, however, was not specific to the stimulation site (i.e., it likewise occurred for saccades to the stimulated and unstimulated side), suggesting a general alerting rather than stimulation specific effect. Landing precision was not affected by V1/V2 or rFEF+ TMS at any tested timepoint.

### Visual areas V1/V2 causally modulate presaccadic benefits shortly before saccade onset (Exp.2a)

To investigate the causal role of early visual areas on the effects of presaccadic attention throughout saccade preparation, we evaluated visual sensitivity at the saccade target–where presaccadic attention *benefits* performance– and opposite of the saccade target–where presaccadic attention *impairs* performance– as a function of V1/V2 stimulation time relative to saccade onset (*Methods – Quantification & statistical analysis*). Given that the saccade target either matched the V1/V2 stimulated region (**Fig. 1c**; *Methods – Phosphene mapping & stimulus placement*) or the ‘symmetric region’ unaffected by TMS (internal control), we can directly assess the effect of V1/V2 TMS on presaccadic attention by comparing visual sensitivity (d’) between these two conditions.

We conducted a repeated measures ANOVA with the factors presaccadic attention (valid/invalid), TMS site (test stimulated/not stimulated), and TMS time (175±25ms/125±25ms/75±25ms/25±25ms prior to saccade onset). Visual sensitivity was higher in valid than invalid trials (*F*(1,9)=6.44, *p*=0.035). Main effects of TMS site (*F*<1) and TMS time (*F*<1) were not significant, and neither were any of the two-way interactions (all *p*>.09). However, a significant 3-way interaction (*F*(3,27)=3.89, *p*=0.026) emerged because V1/V2 TMS significantly reduced presaccadic benefits at the saccade target (compared to when the saccade target was not stimulated) when applied within the last 50ms prior to saccade onset (*p*=0.008, *d*=1.08 **Fig. 3a**–left). Thus, early visual areas become crucial for presaccadic benefits in the final stage of saccade programming, shortly before saccade onset – when presaccadic attention reaches its maximum effect^4–6,72–75^. Presaccadic costs opposite the saccade target (**Fig. 3a**–right) were not affected by V1/V2 TMS at any timepoint. A repeated measures ANOVA [presaccadic attention (valid/invalid) * TMS time] showed that within 50ms before saccade onset, the effect of V1/V2 TMS (computed as test stimulated–not stimulated; **Fig. 3b**) on sensitivity at the saccade target (valid) was significantly stronger than opposite of it (invalid) (attention*TMS_time: *F*(3,27)=3.89, *p*=0.021; valid vs. invalid at 25+-25ms: *p*=0.047, *d*=0.51) – documenting that early visual areas play a selective causal role for presaccadic benefits shortly before saccade onset.

**Figure 3.**
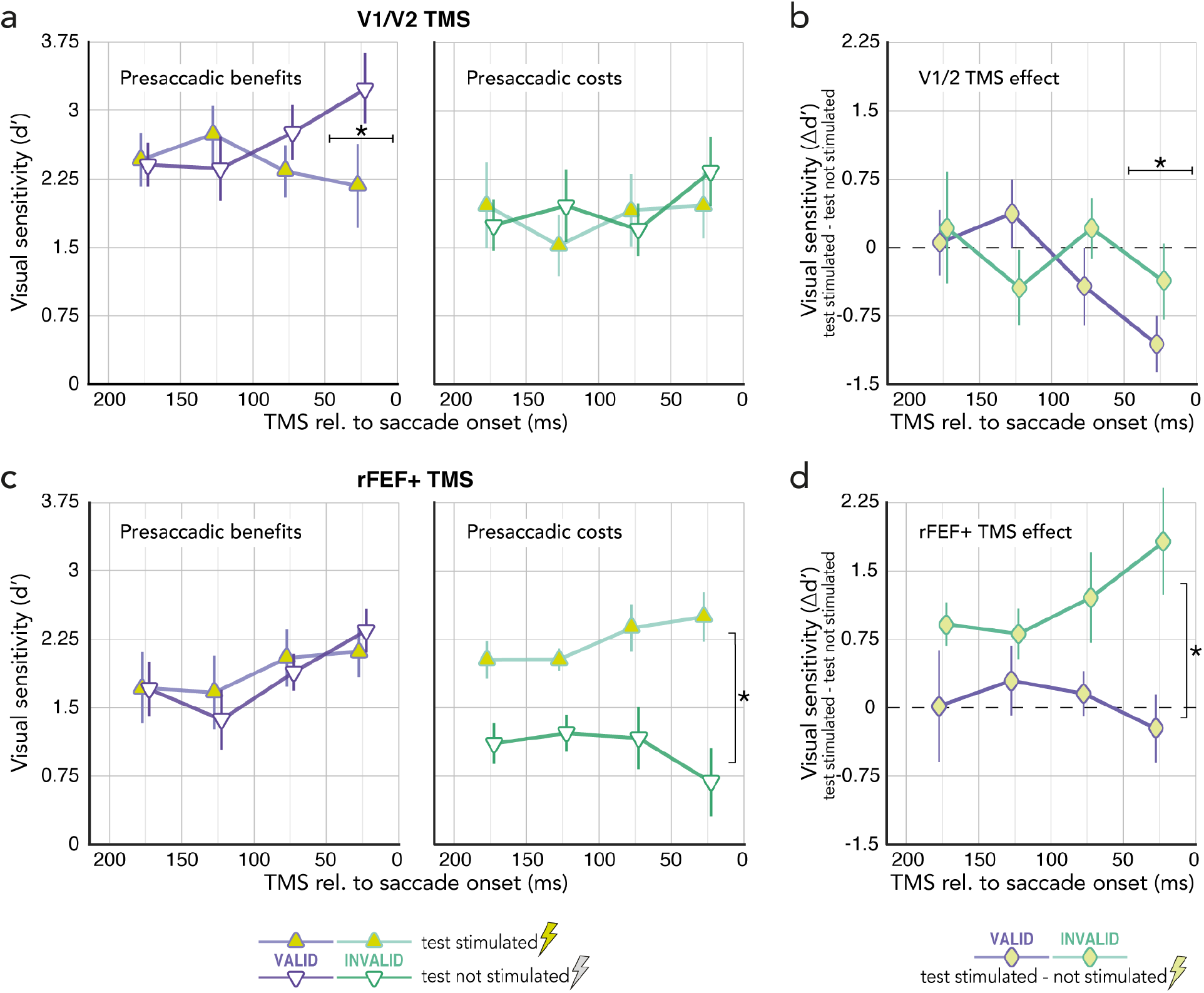
The effect of occipital TMS (V1/V2, **Exp.2a**; upper row) and frontal TMS (rFEF+, **Exp.2b**; lower row) on presaccadic benefits and costs throughout saccade preparation. (**a**) Visual sensitivity at the saccade target (purple) and opposite the saccade target (green) measured at region matching V1/V2 stimulation (yellow triangles) or opposite of it (white triangles, control), binned as a function of stimulation time relative to saccade onset. (**b**) The effect of V1/V2 TMS on presaccadic *benefits* at the saccade target (purple; *valid* test stimulated–not stimulated) and *costs* opposite the saccade target (green; *invalid* test stimulated–not stimulated) binned as a function of stimulation time relative to saccade onset. (**c**) The effect of rFEF+ TMS on visual sensitivity at the saccade target and opposite of it across time; conventions as in (a). (**d**) The effect of rFEF+ TMS on presaccadic benefits and costs across time; conventions as in (b). All symbols and error bars represent the group average ±1SEM. Asterisks indicate significant differences between the two compared conditions at a respective time point (a,b; post-hoc comparison after significant 3-way interaction) or between the two compared conditions across time (c,d; significant main effect).

### rFEF+ stimulation reduces presaccadic costs throughout saccade preparation (Exp.2b)

To assess the contribution of human frontal eye fields to the effects of presaccadic attention throughout saccade preparation, we stimulated rFEF+ and observed a different pattern from that of V1/V2 stimulation, using the otherwise identical experimental protocol, analysis and participant sample (though 3 observers were no longer available). The repeated measures ANOVA [presaccadic attention (benefits/costs) * TMS_site (test stimulated/not stimulated) * TMS_time (175±25ms to 25±25ms before saccade onset) showed no significant main effect of attention (*F*(1,6)=1.11, *p*=0.332) or stimulation time (*F*(3,18)=2.65, *p*=0.110), but a significant main effect of TMS site (*F*(1,6)=12.90, *p*=0.011). The TMS site also interacted with presaccadic attention (*F*(1,6)=8.23, *p*=0.028), which is explained by increased visual sensitivity at the (stimulated) location opposite the saccade target caused by rFEF+ stimulation throughout saccade preparation, i.e., rFEF+ TMS reduced presaccadic costs; *p*=0.012, *d*=1.35; **Fig. 3c**–right). In contrast, rFEF+ TMS did not affect sensitivity at the (stimulated) saccade target (*p*=0.408; **Fig. 3c**–left). Neither the 3-way interaction was significant (*F*(3,18)=1.44, *p*=0.269), nor did presaccadic attention (*F*(3,18)=1.76, *p*=0.200) or stimulation site (*F*(3,18)<1) interact with stimulation time. A repeated measures ANOVA [presaccadic attention (valid/invalid) * TMS time] on the effect of rFEF+ TMS (again computed as test stimulated–not stimulated; **Fig. 3d**) showed a main effect of presaccadic attention (*F*(1,6)=8.23, *p=*0.028); the main effect of stimulation time (*F*<1) and its interaction with presaccadic attention (*F*(3,18)=1.44; *p=*0.268) were not significant. The effect of rFEF+ stimulation on presaccadic costs (enhancing sensitivity opposite the saccade target) thus was stronger than on presaccadic benefits at the saccade target, independent of stimulation time.

## Discussion

This is the first saccade study to investigate the effects of V1/V2 and FEF+ TMS using the same experimental design and stimulation protocol, which enables us to compare the causal role of these brain regions in presaccadic attention. In two psychophysics-TMS experiments we dissociated the involvement of early visual cortex (V1/V2) in presaccadic and covert exogenous^50^ and endogenous attention^63^ (deployed in the absence of eye movements), and document the differential role of frontal (rFEF+) and visual areas (V1/V2) as well as their interplay in presaccadic attention.

V1/V2 TMS did not affect presaccadic attention – unlike covert exogenous attention, which is extinguished^50^ with the same psychophysics protocol and TMS pulse timing (**Fig. 2** & **Fig. S1**). This finding causally dissociates the two forms of attentional orienting, providing further evidence against the claim^77–79^ that covert *exogenous* attention is functionally equivalent to oculomotor programming. The observed effects of presaccadic attention seem similar to those of covert *endogenous* attention^63^, in that they both are affected by FEF+ stimulation. However, perceptual benefits and costs of presaccadic attention are stronger than those of covert endogenous (and exogenous) attention (**Fig. 2b**). Moreover, rFEF+ TMS reduced both the benefits and costs of endogenous attention^63^, but it only reduced presaccadic costs at the non-target location, leaving (the typically high) presaccadic sensitivity at the saccade target unaffected (**Fig. 3c**).

The rFEF+ TMS induced enhancement opposite the saccade target is consistent with evidence that TMS effects depend on the brain activation state at the time of stimulation^50,63,80–86^: TMS is more likely to increase performance where it is usually worse (e.g., opposite the saccade target). It is also in line with the view of FEF being a priority map, controlling / shifting the focus of attention focus^29,30,87^. Similarly, FEF-microsimulation in non-human primates increases visual sensitivity at the movement field of the stimulated neurons^35–37^, akin to a shift of attention.

Previous TMS-studies investigating the influence of FEF+ on presaccadic attention have yielded mixed results, reporting that FEF+ TMS either decreased^88^ or increased^89^ sensitivity at the saccade target. Crucially, these studies have not evaluated the TMS-effect relative to saccade onset (as we do in Experiment 2). However, the effects of presaccadic attention gradually increase throughout saccade preparation^13^ and saccadic reaction times vary profoundly between (and within) individual observers – which is why presaccadic attention effects typically are evaluated relative to saccade onset^4,73–75^. The inconsistent results of previous TMS studies^88,89^ could be explained by FEF+ stimulation being applied at different time points relative to saccade onset (i.e., when the effects of presaccadic attention would be differently pronounced).

V1/V2 TMS shortly (within 50ms) before saccade onset, i.e., at the peak of presaccadic attention, reduced the typical sensitivity benefit at the saccade target. Importantly, this effect was time-locked to saccade onset, indicating that occipital regions are recruited shortly before the eyes move. Applied at a fixed time relatively earlier during saccade programming (100ms after cue onset), V1/V2 TMS did not affect presaccadic attention.

To conclude, our results dissociate the neural basis of presaccadic from covert attention and demonstrate a causal and differential role of occipital and frontal areas in presaccadic benefits (at the saccade target) and costs (opposite of it). Whereas the effect of frontal TMS was present throughout saccade preparation, critically, the effect of occipital TMS was locked to the period right before saccade onset, which – consistent with presaccadic feedback from oculomotor structures to visual cortex^29,30,36,90^ – reveals that occipital regions are recruited only during later stages of saccade programming.

## Additional information and files

### Author contributions

Conceptualization: NMH and MC; Methodology and software: NMH; Investigation: NMH; Formal analysis: NMH and AF; Visualization: NMH; Writing – original draft: NMH; Writing – review & editing: NMH, AF, and MC; Funding acquisition: NMH and MC.

### Data availability

Raw eye tracking and behavioral data will be available from the OSF database URL: https://osf.io/pcunw/.

### Supplemental Video S1

Demonstration of the trials sequence. Shown are one *valid*, one *invalid*, and one *neutral* trial.

## Acknowledgements

This research was supported by a Marie Skłodowska-Curie individual fellowship (MSCA-IF 898520) by the European Commission to NMH, an NIH NINDS grant (F99-NS-120705) to AF, and an NIH NEI grant (R01-EY-019693) to MC. We thank the Carrasco Lab members, in particular Yuna Kwak and Hsing-Hao Lee, as well as Ilona Bloem and Jan Kurzawski for helpful comments and discussions.

## Methods

### Observers

10 observers (7 female; aged 22–36 years) participated in Experiment 1. 10 observers (8 of which participated in Experiment 1; 8 female; aged 21–36 years) participated in Experiment 2a, of which 7 observers (7 female; aged 22–36 years) also participated in Experiment 2b. All observers had normal or corrected-to-normal vision, provided written informed consent, and (except for one author) were naive to the purpose of the experiment. We chose a sample size in the range of previous psychophysics-TMS studies investigating presaccadic and covert attention^50,63,88,89^. Observers were screened for TMS counterindications prior to participation. The protocols for the study were in accordance with the safety guidelines for TMS research and approved by the University Committee on Activities Involving Human Subjects at New York University and all experimental procedures were in agreement with the Declaration of Helsinki.

### Setup

Observers sat in a dark room with their head stabilized by a chin and forehead rest and viewed the stimuli at 57 cm distance on a gamma-linearized ViewPixx/EEG LCD monitor (VPixx Technologies, Saint-Bruno, QC, Canada) with a spatial resolution of 1,920 by 1,080 pixels and a vertical refresh rate of 120 Hz. Gaze position of the dominant eye was recorded using an EyeLink 1000 Desktop Mount eye tracker (SR Research, Osgoode, Ontario, Canada) at a sampling rate of 1 kHz. Manual responses were recorded via a standard keyboard. A Linux desktop machine running Matlab (MathWorks, Natick, MA, USA) with Psychophysics^91,92^ and EyeLink toolboxes^93^ controlled stimulus presentation and response collection.

### Phosphene mapping & stimulus placement

Prior to Experiment 1, observers fixated a fixation target at the center of the black screen. We applied a train of seven TMS pulses (30 Hz, 65% of maximal stimulator output) at the assumed phosphene region (laterally around the occipital pole). Once the observer perceived a reliable phosphene in the contralateral visual field, they drew its outline on the screen using a computer mouse, and the exact TMS coil position and angle was recorded using the Brainsight TMS navigation system (Rogue Research, Montréal, QC, Canada). The center of the phosphene drawing was used for stimulus placement in Experiment 1 and 2 (see **Fig. 1c**) – where we stimulated the same region, but with verified sub-threshold stimulation intensity (i.e., observers did not perceive phosphenes during the main experiments), to not contaminate the measure and capture attention ‘visually’^94^. TMS coil positions eliciting phosphenes were validated before each experimental session. In Experiment 1, 7 observers perceived phosphenes in the right visual field and 3 observers perceived phosphenes in the left visual field (average eccentricity from fixation: 6.39°±0.63°; mean±1SEM). In Experiment 2, 6 observers perceived phosphenes in the right visual field and 4 observers perceived phosphenes in the left visual field (average eccentricity 6.68°±0.66°). For the analysis of behavioral results, we collapse data across phosphene sides, as previous studies stimulating V1/V2 have found no side-specific effects^50,63,70^.

### FEF Localization

Before participating in Experiment 2a, we localized each observers (human) right Frontal Eye Field (rFEF+) using the Wang atlas^76^, which has been shown to be a reliable indicator of FEF+^63,95,96^. We mapped the right FEF+ onto each observer’s native volume using mri_surf2vol & mri_surf2surf in Freesurfer^97^ (**Fig. 1d**). The rFEF+ region of interest (ROI) was validated via anatomical landmarks – the junction of the precentral and superior frontal sulci^64,65,67^. Observer’s individual anatomical brain scan (T1 image) and rFEF+ ROI were loaded into the neuro-navigation software Brainsight (Rogue Research, Montréal, QC, Canada) for precise stimulation of rFEF+ in Experiment 2b.

### Experimental design

**Experiment 1** (see **Fig. 1e** & **Video S1**). Observers fixated a central fixation target comprising a black (~0cd/m2) and white (~96cd/m2) bull’s-eye (r=0.3°) on gray background (~48cd/m2). Two placeholders indicated the two potential saccade target locations left and right of fixation, each comprised four black dots (r=0.15°). Saccade target centers were determined for each observer via phosphene mapping (see *Phosphene mapping & stimulus placement*). Once stable fixation was detected within a 1.75° from fixation for at least 250 ms, the trial started with a jittered fixation period (250ms, 750ms, or 1,250ms), before a central direction cue (black line, length 0.75°) pointed to one of the placeholders, thereby cueing the saccade target. Observers were instructed to “look as fast and precisely as possible” to the center of the indicated placeholder. Note that the saccade was equally likely directed to the *stimulated* phosphene region and to the non-stimulated *symmetric* region (**Fig. 1c**). 100ms after saccade cue onset (i.e., within the movement latency, gaze still rests at fixation), Gabor gratings (2cpd, random phase) appeared for 100ms within each placeholder; one grating was vertical, the other *test* grating was slightly tilted (see *Titration procedure*). Gabor contrast varied from trial to trial (method of constant stimuli). Gabor size was adjusted according to the Cortical Magnification Factor: [M = M0(1+0.42E+0.000055E3)-1]^98^, where M0 refers to the cortical magnification factor (7.99 mm/deg) and E to the stimulus eccentricity in degrees of visual angle; Gabors were scaled to match a cortical magnification of a 2° wide Gabor at 4° eccentricity. Placeholder dots were separated by 1° from the Gabors. The first TMS pulse was time-locked to Gabor onset, followed by another pulse 50ms later. 400ms after Gabor offset (the eye movement has now been performed), the dots of one placeholder changed color from black to white, functioning as a response cue to indicate the location that had contained the tilted Gabor. Observers indicated their orientation judgement via button press (clockwise or counterclockwise, two-alternative forced choice) and were informed that the orientation report was non-speeded. They received auditory feedback for incorrect responses. Importantly, the tilted test Gabor was equally likely presented at the saccade target (*valid* trials) or at the opposite, non-target location (*invalid* trials), i.e. the saccade cue was not predictive of the test location, valid and invalid trials were randomly intermixed. In separately blocked *neutral* trials, two central line cues indicated both placeholder locations and participants were instructed to keep fixating. After an initial training without TMS, observers performed 18 experimental blocks (12 saccade blocks with valid and invalid trials, 6 fixation blocks with neutral trials); random order, 160 trials per block) split into 3 experimental sessions. We controlled online for broken eye fixation (outside 1.75° from central fixation before the cue onset), too short (<150ms) or too long (>500ms) eye movement latencies, and imprecise saccades (not landing within 2.5° from saccade target center). Erroneous trials were repeated in random order at the end of each block.

In **Experiment 2** we used the same psychophysics-TMS as in Experiment 1, with the following differences: (*1*) Double-pulse V1/V2 (Exp. 2a) or rFEF+ (Exp. 2b; see *FEF Localization*) TMS was applied at different time points throughout saccade preparation (first TMS pulse 0-200ms relative to cue onset; **Fig. 1f**). (*2*) Whereas grating contrast in Experiment 1 was varied to measure contrast response functions, the contrast was fixed to 46% in Experiment 2. (*3*) We only tested *valid* and *invalid* trials (randomly intermixed), with separately determined Gabor tilt angles (see *Titration procedure*). Note that stimulus placement was individually determined for each observer (see *Phosphene mapping & stimulus placement*), but identical for both Exp. 2a and 2b. Observers performed 7 (Exp. 2a) or 10 (Exp. 2a) experimental blocks of 160 trials each, split into 2 experimental sessions. We again repeated erroneous trials with broken eye fixation (outside 1.75° from central fixation before the cue onset), too short (<150ms) or too long (>500ms) eye movement latencies, or imprecise saccades landing further than 2.5° from saccade target center.

In total, we included 22,679 trials in the analysis of the behavioral results of Experiment 1 (on average 2,268 ± 35 (mean±1SEM) trials per observer), 5,295 (530±13) trials in the analysis of Experiment 2a, and 5,868 (838±31) trials in the analysis of Experiment 2b.

### Titration procedure

To match overall task difficulty to each observers ’visual sensitivity and to account for any learning effects, we titrated the Gabor-tilt angle ( ±0.5°–6° relative to vertical) separately for each observer before each experimental session (without TMS) via an adaptive staircase procedure^99^ implemented in the Palamedes toolbox^100^. For Experiment 1, we used the procedure of the *neutral* condition to determine the tilt angle at which observers ’orientation discrimination performance at the highest contrast level (85%) was ~d’ = 2. This tilt angle (group average 1.7±°0.2°) was used in the main experiment for the *valid, invalid*, and *neutral* conditions. For Experiment 2, to account for the observed pronounced sensitivity differences at and opposite the saccade target, we the tilt angle during saccade preparation separately for *valid* (Exp. 2a: 1.1±°0.1°; Exp. 2b: 1.1±°0.2°) and *invalid* (Exp. 2a: 4.1±°1.0°; Exp. 2b: 4.O±°1.5°) trials using a Gabor contrast of 46% (same contrast as in main experiment).

### Eye data preprocessing

We scanned the recorded eye-position data offline and detected saccades based on their velocity distribution^101^ using a moving average over 20 subsequent eye position samples. Saccade onset and offset were detected when the velocity exceeded or fell below the median of the moving average by 3 standard deviations for at least 20ms. We included trials in which no blink occurred during the trial and correct eye fixation was maintained within a 1.75° radius centered on central fixation throughout the trial (fixation trials) or until cue onset (saccade trials). Moreover, we only included those eye movement trials in which the initial saccade landed within 2.0° from the required target location and in which the test signal was presented within 100ms before saccade onset (i.e., the saccade started only after test signal presentation, but not later than 100ms after signal offset)

### Quantification & statistical analysis

Task performance, indexed by visual sensitivity [d-prime; d ’= *z*(hit rate) - *z*(false alarm rate)], was measured as a function of stimulus contrast using the method of constant stimuli (6 Michelson contrast levels: 2, 7, 13, 24, 46, 85%). We arbitrarily defined counter-clockwise responses to counter-clockwise oriented gratings as hits and counter-clockwise responses to clockwise oriented gratings as false-alarms^9,48,50,63,102^. To avoid infinite values when computing d’, we substituted hit and false alarm rates of 0 and 1 by 0.01 and 0.99, respectively^5,15,16,103^.

To obtain contrast response functions for each condition and test location we fit each observer’s data with Naka-Rushton functions^45^, parameterized as d’(C) = d_max_C^n^ / (C^n^ + C^n^_50_), where C is the contrast level, dmax is the asymptotic performance, C_50_ is the semi-saturation constant (contrast level corresponding to half the asymptotic performance), and n determines the slope of the function. The error was minimized using a least-squared criterion; d_max_ and c_50_ were free parameters, n was fixed. Contrast levels were log-transformed prior to fitting. A change in d_max_ indicates a response gain change.

We used repeated measures ANOVAs to assess statistical significance, followed by Bonferroni-corrected multiple (post-hoc) comparisons, if applicable. For repeated-measures ANOVAs in which the sphericity assumption was not met, we report Greenhouse-Geisser corrected *p*-values.

For Experiment 2, we binned trials as a function of the time between Gabor stimuli offset and saccade onset (see *Eye data preprocessing*) in four separate time windows (200–150ms, 150–100ms, 100–50ms, and 50–0ms).

## Supplemental figures

**Supplemental Figure S1.**
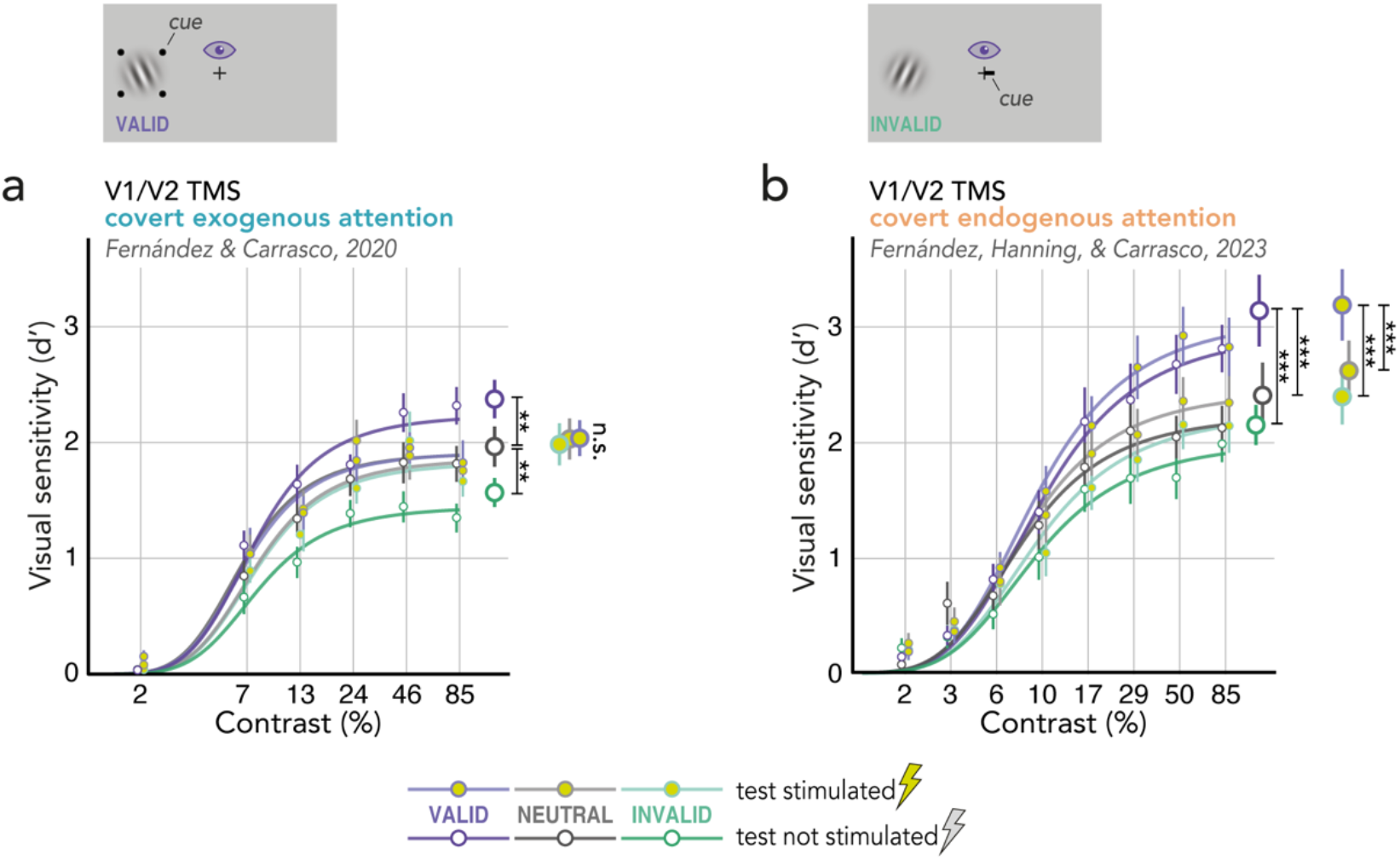
The effect of V1/V2 TMS on covert attention. (**a**) Exogenous attention (data previously published^50^). Contrast Response Functions (CRF) for *valid* trials (target at cued location; purple), *invalid* trials (target opposite cued location; green), and *neutral* trials (both locations cued; gray), measured at the stimulated region (yellow symbols) or in the symmetric, nonstimulated hemifield (white symbols). Respective group averaged parameter estimates for the upper asymptote d_max_ (based individual observers’ fits) displayed on the right. Error bars indicate ±1SEM. (**b**) Endogenous attention (data previously published^63^). Same conventions as in (a). ** *p*<.01, *** *p*<.001

**Supplemental Figure S2.**
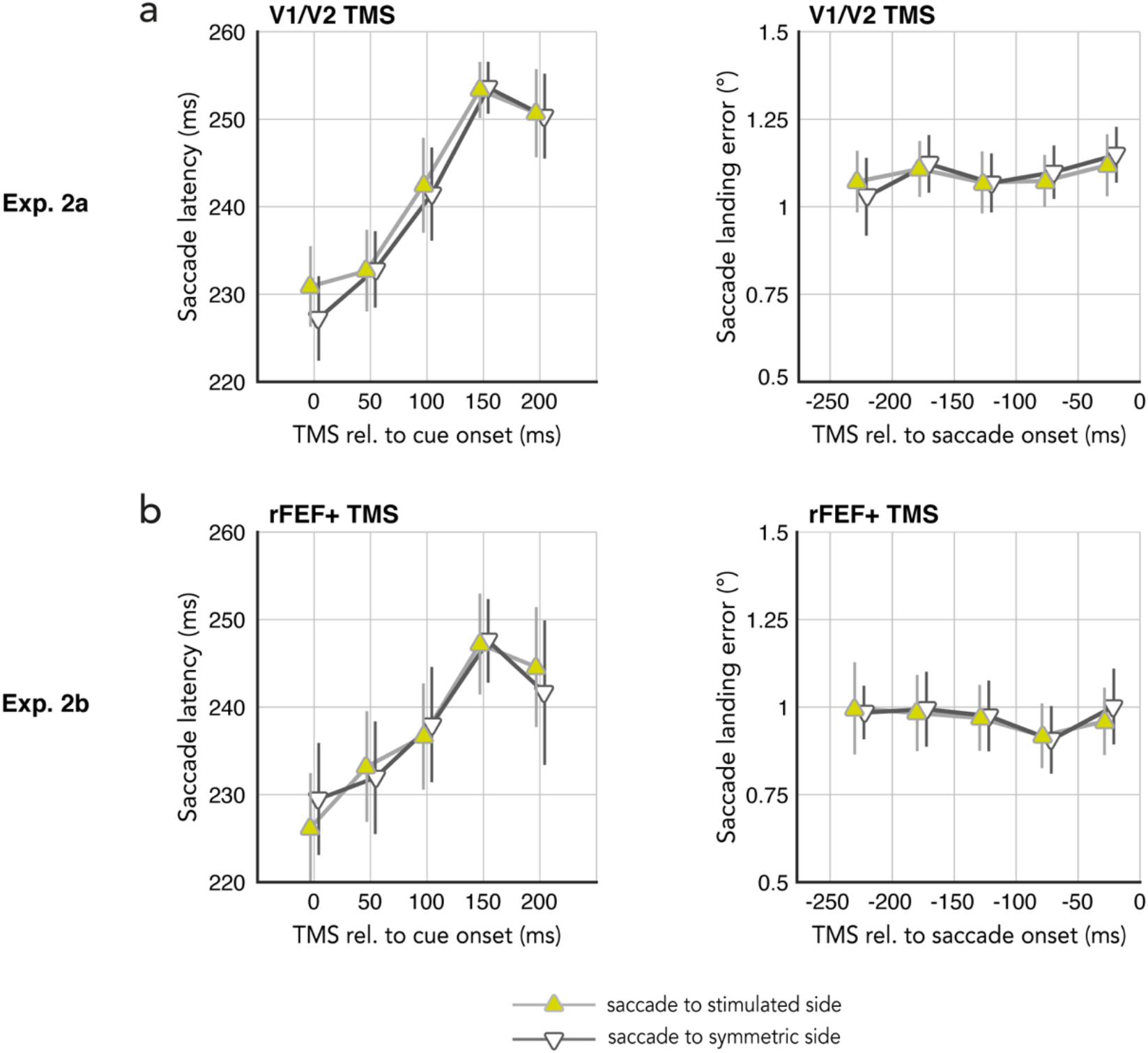
Eye movement parameters. Saccade latencies (left) and precision (landing error, computed as the distance between saccade landing position and saccade target center; right) in Exp.2a, V1/V2 TMS (**a**), and Exp.2b, rFEF+ TMS (**b**). Data plotted as a function of TMS pulse relative to cue onset (saccade latencies, left) or TMS pulse relative to saccade onset (landing error, right). TMS over both stimulation sites increased saccadic latencies (the later the TMS pulse, the later saccade onset). This pattern was not side-specific (it occurred similarly for saccades directed to the stimulated and symmetric side), and thus likely is explained by a general alerting effect of the TMS sound / sensation rather than the stimulation itself. Saccade precision was not affected by V1/V2 or rFEF+ TMS at any tested timepoint.

